# Intracranial electrical stimulation alters meso-scale network integration as a function of network topology

**DOI:** 10.1101/2021.01.16.426941

**Authors:** WH Thompson, O Esteban, H Oya, R Nair, F Eberhardt, J Dubois, RA Poldrack, R Adolphs, JM Shine

## Abstract

Human brain dynamics are organized into a multi-scale network structure that contains multiple tight-knit, meso-scale communities. Recent work has demonstrated that many psychological capacities, as well as impairments in cognitive function secondary to damage, can be mapped onto organizing principles at this mesoscopic scale. However, we still don’t know the rules that govern the dynamic interactions between regions that are constrained by the topology of the broader network. In this preregistered study, we utilized a unique human dataset in which whole brain BOLD-fMRI activity was recorded simultaneously with intracranial electrical stimulation, to characterize the effects of direct neural stimulation on the dynamic reconfiguration of the broader network. Direct neural stimulation increased the extent to which the stimulation site’s own mesoscale community integrated with the rest of the brain. Further, we found that these network changes depended on the topological role of the stimulation site itself: stimulating regions with high participation coefficients led to global integration, whereas stimulating sites with low participation coefficients integrated that regions’ own community with the rest of the brain. These findings provide direct causal evidence for how network topology shapes and constrains inter-regional coordination, and suggest applications for targeted therapeutic interventions in patients with deep-brain stimulation.

## Introduction

Cognition emerges from flexible, dynamic interactions between neurons distributed across the central nervous system. Advances in systems neuroscience have demonstrated a number of analytical tools for tracking these coordinated dynamics over time in whole brain imaging data. For instance, dimensionality reduction approaches can be used to summarize network-wide activity using a much smaller number of components (Cunningham & Yu, 2014), whose activity can then be tracked over time (Gallego et al., 2017, 2020). Similarly, the macroscopic network organization of the brain has been shown to fluctuate between periods of integration and segregation (Betzel et al., 2016; Shine et al., 2016). Despite promising links to behaviour, the neuroscientific interpretation of these fluctuations however remains an open scientific question (Shine & Poldrack, 2018). There is evidence to suggest that the brain can shift towards integration during task performance (Cohen & D’Esposito, 2016; Shine et al., 2016), while others have argued that the interplay of a subset of segregated systems is also crucial (Fransson et al., 2018). These lines of inquiry suggest that the dynamic balance between integration and segregation is perhaps the most critical feature of whole-brain organization (Park & Friston, 2013). However, we still do not know the rules that govern transitions between meso-scale topological states in the awake, human brain.

Obtaining causal evidence for system-wide interactions in the human brain is inherently challenging. Even intracranial EEG, which affords direct access to the brain, is associated with limited source localization and minimal access to the dynamics of the brain that are outside the recording site. Deep brain and cortical stimulation have revealed that network changes occur following stimulation (Alhourani et al., 2015; Huang et al., 2019; Shine et al., 2017), however the whole-brain network changes resulting from intracranial electrical stimulation are often difficult to identify, due to the sparsity of sampling points in space (i.e., cortical coverage is typically only partial) and that stimulation often occurs outside the magnetic resonance imaging (MRI) scanner. On the other hand, blood-oxygen level-dependent (BOLD) fMRI can be collected from the whole brain, but it is challenging to causally perturb the system in a direct manner, as TMS-fMRI only stimulates cortical tissue indirectly.

Recently, we demonstrated that stimulation of intracranial EEG electrodes can be successfully combined with functional MRI (fMRI) to simultaneously solve these problems (es-fMRI) (Dubois et al., 2017; Oya et al., 2017; Thompson et al., 2020). In previous work, we showed that there are consistent BOLD fMRI patterns that arise following focal electrical stimulation through depth electrodes. Here we leveraged this dataset to investigate how experimental intracranial stimulation at specific network nodes would influence network structure at the whole-brain level. In a pre-registered analysis (https://osf.io/pdhfu/; including seven explicit deviations, which are detailed in the Supplementary Materials), we analysed data from 26 subjects with epilepsy who underwent es-fMRI (Thompson et al., 2020). We set out to investigate two related questions. Our first research question (RQ1) tested whether nodes in the stimulation site’s community (compared to nodes in other communities) caused a reconfiguration of network topology, specifically the balance between network integration and segregation. Our second research question (RQ2) asked whether stimulation-driven changes in network properties were dependent on the topological role of the stimulation site. In both cases, we observed predicted effects that provide insights into global network effects caused by direct electrical stimulation in awake human participants. Together, these results help to define the principles that govern the coordinated interactions between regional- and network-level organization in the human brain.

## Results

The primary focus of our study was on the difference between the periods with and without electrical stimulation (es-on/off, respectively) (Figure 1A). To quantify regional signatures of network topology, we utilized two summary metrics: participation coefficient (PC) and module degree z-score (z), which are together sometimes referred to as a cartographic profile (Guimera et al., 2005). The participation coefficient (PC) quantifies the distribution of network edges (Figure 1B), and indicates how a node in a given community is linked with different modules across the network. In contrast, the module degree z-score (z) quantifies the standardized strength of connections an individual node has within its own community, relative to the mean strength of the module (Figure 1B). Together, these two metrics quantify the integration and segregation of nodes within the context of the whole network: high PC is indicative of relative integration, whereas high z is reflective of relative segregation (Guimera et al., 2005; Power et al., 2011). For each subject/run combination, we quantified the difference between es-on and es-off in PC and z. These es-on/off differences were computed for all nodes *within* a stimulation site’s community and for all nodes *outside* the stimulation site’s community, yielding four measures of interest: ΔPC_within_, ΔPC_outside_, Δz_within_, and Δz_outside_, where the ‘within/outside’ subscript indicates the relationship with the stimulation site’s community, Δ denotes the difference between es-on and es-off, and PC/z.

**Figure 1:**
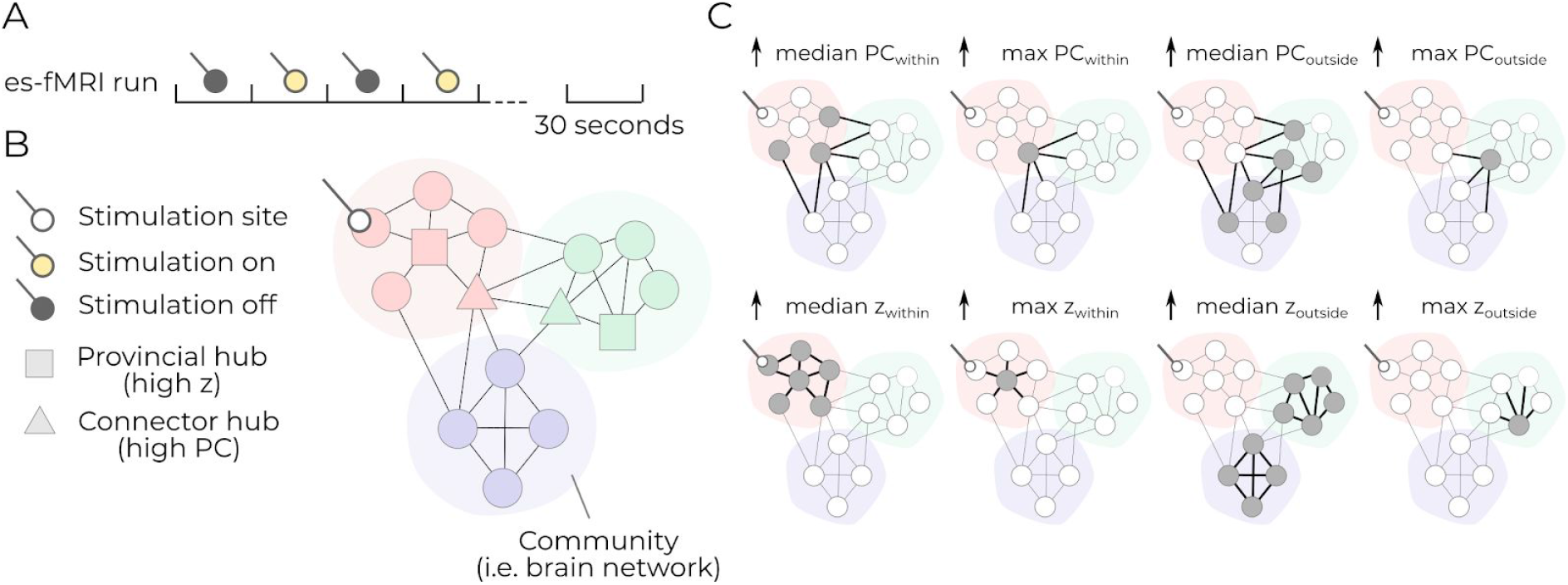
Conceptual figure outlining the network level analyses used in the article. **A**. An example of the block design of the es-fMRI run where, during each run, the electrical stimulation pulses were on for 30 second periods. See (Thompson et al., 2020) for details. **B**. An schematic network illustrating nodes with high participation coefficient (PC) and high within module degree z-score. An example stimulation site is shown in the red community. **C**. An illustration of where changes would occur in the network for an *increase* in each of the 8 summary measures used throughout this paper. The gray nodes show which nodes are affected by the increase. The thicker black lines show the edges that would be affected. Note the distinction between “within” and “outside” the stimulation site community, and the distinction at the network-level of network wide changes (median) and single hub changes (max).

Two different summary metrics were used to summarize the overall change following electrical stimulation: the median and the maximum of the nodal properties, both of which have slightly different interpretations (Figure 1C). If we expect that the most prominent nodes, sometimes called ‘hubs’ (van den Heuvel & Sporns, 2013), are the ones primarily affected by the electrical stimulation, there should be an increase in the max PC (particularly for connector hubs, which are nodes with the most diverse connections between communities) and in the max z (for provincial hubs, i.e. those with the most connections within communities). Alternatively, if we expect that electrical stimulation leads to widespread changes in the topological properties of a community (rather than at only the most prominent nodes), then this will instead be reflected in the median PC (if the community, as a whole, integrates more extensively), and median z (if the community, as a whole, segregates more extensively).

### Electrical stimulation induces community-wide integration for the stimulation site’s community (RQ1)

We quantified, for each subject, each of our 8 metrics during those epochs where electrical stimulation was on, and those epochs where electrical stimulation was off. This left us with a total of 16 initial distributions across subjects, whose details are shown in Figure 2A-H. From these distributions, some runs showed an increase when es-on, notably, median PC_within_ (Figure 2A), median PC_outside_ (Figure 2B), and max z_outside_ (Figure 2H), while the median z_outside_ appears to decrease compared when for the es-on periods (Figure 2F). The remaining four summary statistics had approximately 50% of the es-on runs being higher than the es-off runs (46.3 − 51.2%). From these 16 distributions, we derived 8 difference metrics between the on and off periods to test RQ1.

**Figure 2:**
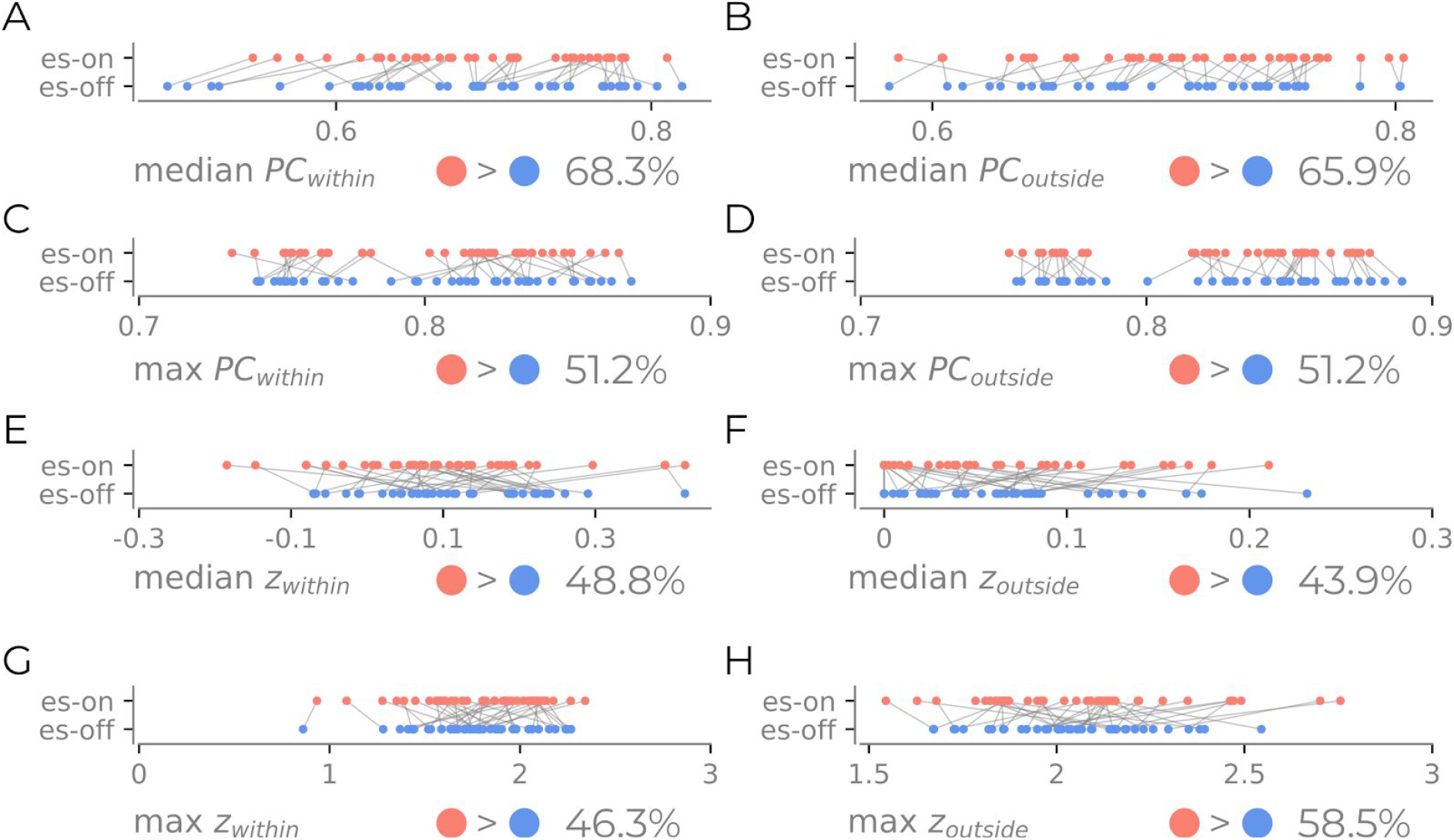
Distribution of 8 network summary statistics per fun for the es-on and es-off periods. **A**. median PC_within_; **B**. median PC_outside_; **C**. max PC_within_; **D**. max PC_outside_; **E**. median z_within_; **F**. median z_outside_; **G**. max z_within_; **H**. max z_outside_;. Each panel shows the percentage of runs where the es-on run was larger than the es-off run.

We asked whether there was a significant difference in the effects evoked between the stimulation site’s within-community and outside-community metrics (i.e., median ΔPC_within_ vs ΔPC_outside_, four comparisons in total). We saw, for example, that in Figure 2 both PC_within_ and PC_outside_ were associated with higher values when the stimulation was on. However, now we ask whether the magnitude of change is greater within or outside the stimulation site’s community. An increase in ΔPC_within_ would indicate whether the topological change caused by electrical stimulation is primarily affecting the stimulation site’s community.

Alternatively, if there is no difference, then electrical stimulation would be affecting global topological properties. The results are shown in Figure 3A-D. In one instance, there was evidence to reject the null hypothesis, namely for the median participation coefficient (avg. difference: 0.013, p=0.0045, Figure 3B). When correcting for the four statistical tests performed with a Bonferroni correction and a significance threshold of 0.05, the median participation can be classed as significantly differing for the stimulation site’s community. For the other measures, the null hypothesis could not be rejected (max ΔPC_within_ vs ΔPC_outside_: 0.0013, p=0.29, Figure 3A; max Δz_within_ vs Δz_outside_: 0.0028, p=0.52, Figure 3C; Median ΔPC_within_ vs ΔPC_outside_: 0.013, p=0.70, Figure 3D). However, it can be observed that both the max ΔPC_outside_and the median Δz_outside_ both decrease tend towards 0 compared to the within the stimulation community, suggesting that these properties can both increase and decrease following electrical stimulation.

**Figure 3:**
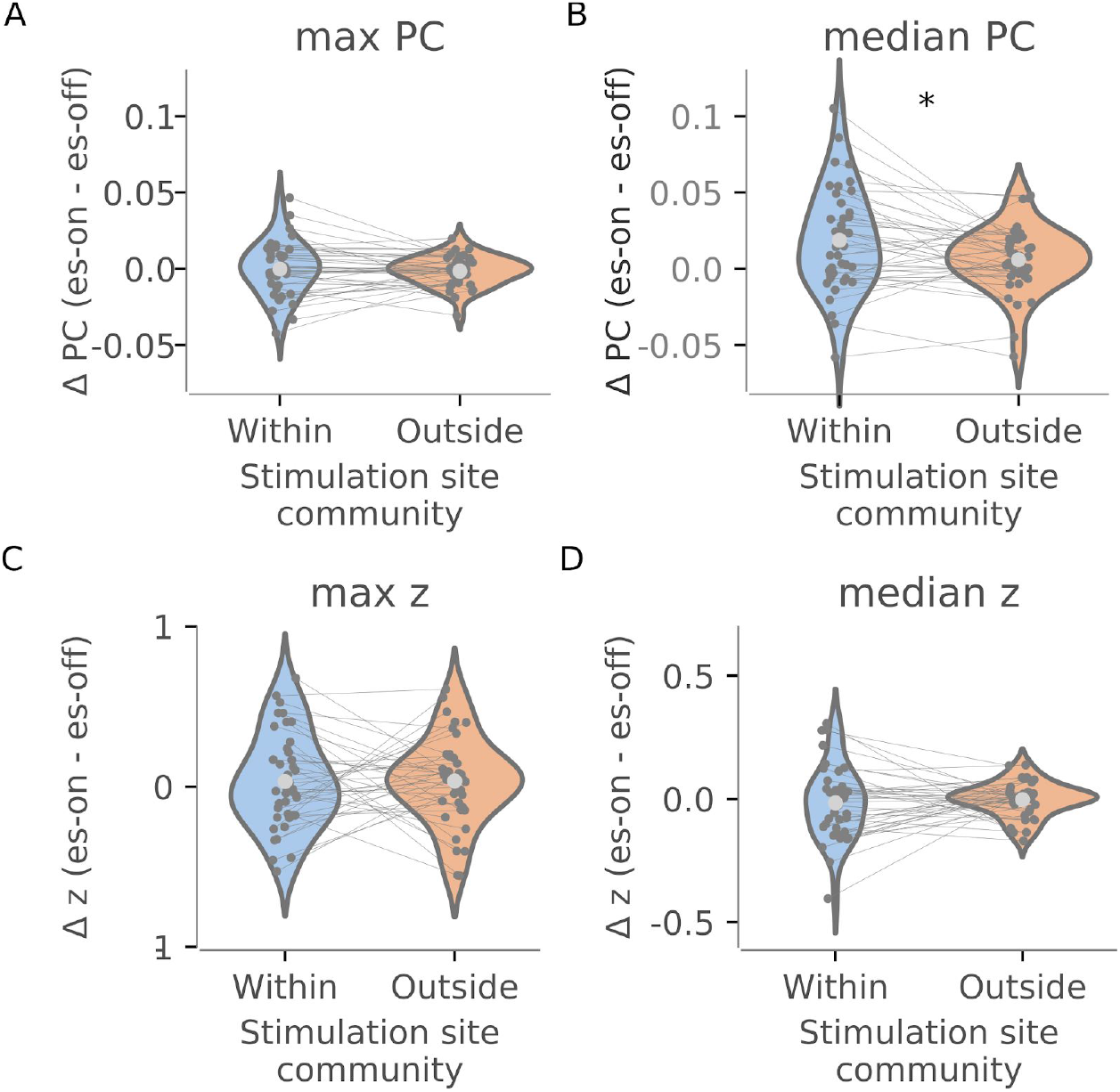
Differences in network properties following stimulation between the stimulation sites community and all other communities. **A**. max ΔPC; **B**, median ΔPC; **C**. max Δz; **D**. median Δz. Light gray dots indicate the average. Each dark gray dot marks a subject’s run, and the lines connect the same run for both measures. * marks p<0.05, Bonferroni corrected.

Finally, we demonstrate that the interpretation of the summary statistics was correct, especially that the median ΔPC_within_ was indeed identifying a general increase in the integration community of all nodes in the stimulation site’s community. Figure 4AB shows the node’s PC and z during es-on vs es-off for a single subject in order to verify this interpretation (Figure 4A/B). In Figure 4A, an ostensible pattern appears, wherein the stimulation site’s community has a general increase in the participation coefficient for most nodes in the es-runs. This shift is captured by the median, illustrating that this measure captures wide-spread changes across the network. In Figure 4B, no clear overall pattern can be seen for all nodes. Together, these results help to justify our interpretation of what is occurring on the nodal level with the community-level summary statistics.

**Figure 4.**
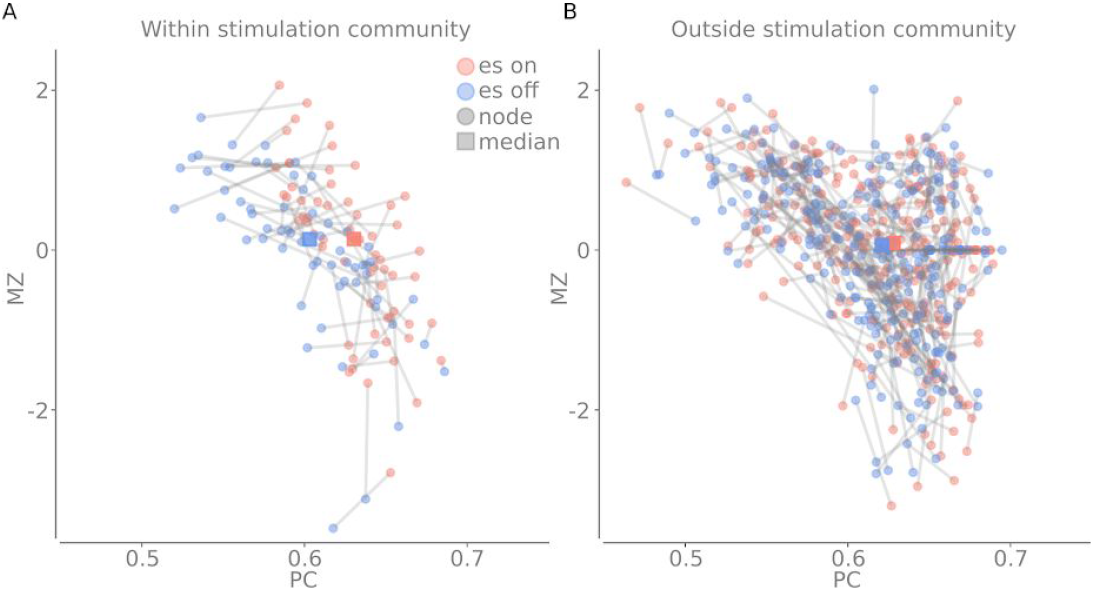
Cartographic profile from a randomly chosen subject/run. **A**. The participation coefficient and within module degree z-score for one subject/run. Each pair of connected points indicate a node from the stimulation site’s community when es-on and es-off. The median changes for the participation coefficient. **B**. Same as A but for all nodes outside of the stimulation site.

Together, the results associated with RQ1 demonstrated that the median PC increases when the electrical stimulation is on, and that this increase is significantly larger for PC_within_. This result entails that there is a community-wide increase in integration within the stimulation site’s community following electrical stimulation. Aside from this, no other significant differences were found between the summary statistics for the topological differences between the within stimulation site community and the outside stimulation site community.

### The rise in within-stimulation site participation coefficient reflects an increase in integration with a subset of communities, not global integration

Thus far we have demonstrated that the median participation coefficient increases for the community of the stimulation site following electrical stimulation. However, this increase in integration could be instantiated in one of two possible ways: (i) a *selected increase −* in which a subsection of other communities increase their connectivity with the stimulation community; or (ii) a *global increase* - in which all communities increase their connectivity with the community of the stimulation site. To arbitrate between these possibilities, we conducted an un-registered *post hoc* analysis in which we quantified the median change in edge strength (i.e., the sum of edge weights) between each node in the stimulation community with all other communities. We observed that the distributions from the increased integration from the stimulation site’s community was generally limited to a subsection of communities, not all communities (Supplementary Figure 1) - thus confirming the first possibility.

### The stimulation site’s topological role affects how the network changes (RQ2)

The previous sections identified a general increase in median participation within the stimulation site’s community. However, it did not consider the topological role of the stimulated node - i.e., whether or not the node itself links many other communities. Given the scale-free nature of the structural connectome of the brain, this topological feature could have an important influence on how the stimulation changes the network. In addition, divergences can be seen in the previous sections where, for some runs, the stimulation site’s community does not increase their integration. This suggests that the extent, or even presence, of a widespread network effect of stimulation may be dependent on the topological properties of the area stimulated.

This question was investigated in our second hypothesis (i.e., RQ2). Note that for most subjects, there were multiple stimulation sites that varied in their participation coefficient (Figure 5A), and also a broad distribution of participation across stimulation sites across all subjects. We ran multiple Bayesian models to examine whether the PC of the stimulation site relates to different combinations of the eight delta summary statistics used previously. All possible combinations were run, and the best-fitting model was calculated using leave-one-out (LOO) information criteria (Vehtari et al., 2017). The best-fitting model contained three variables: median ΔPC_within_, median ΔPC_outside_, and max Δz_outside_ (Figure 5BC, see Methods and Supplementary Table 12).

**Figure 5.**
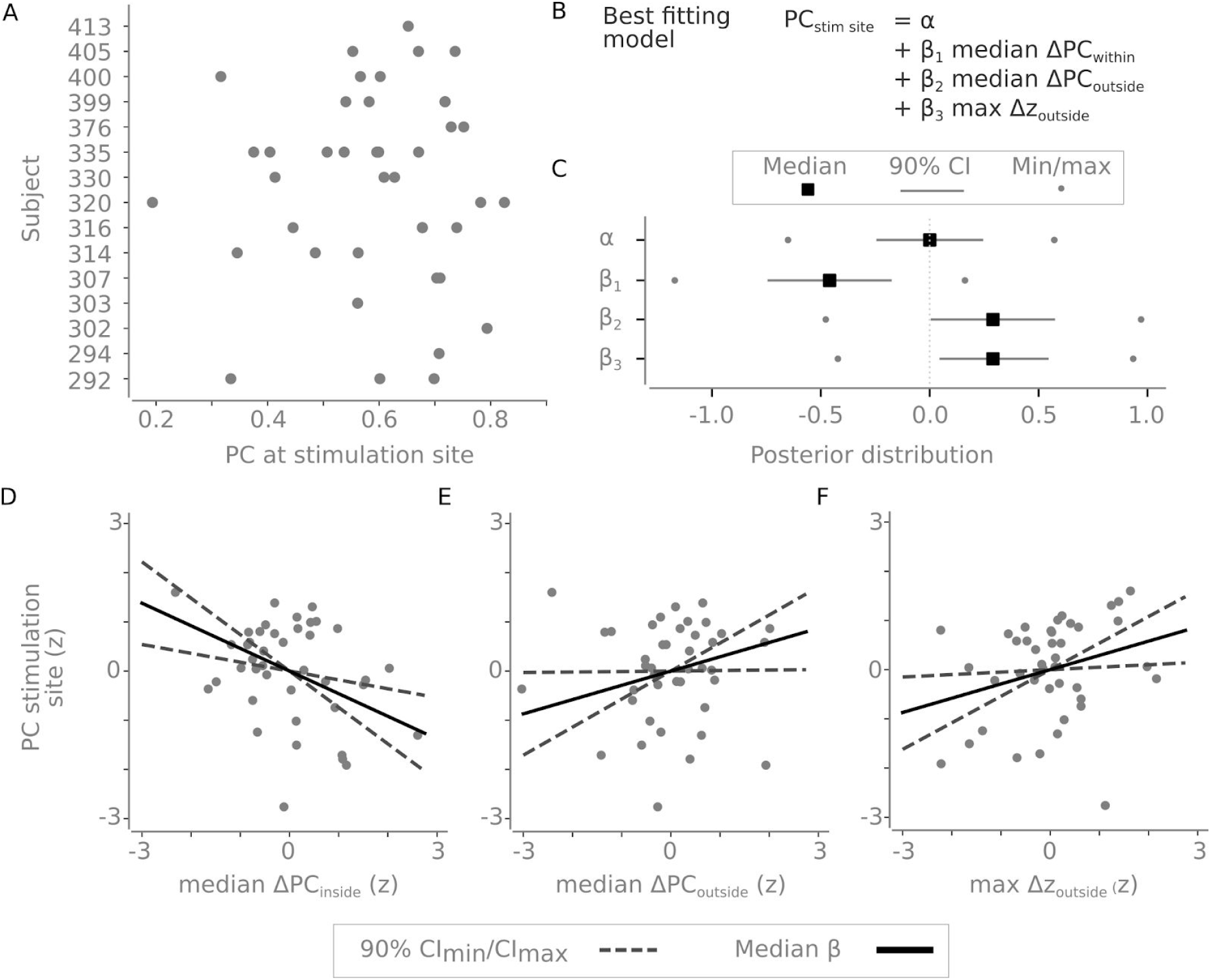
The topological role of the stimulation site relates to network changes. **A**. The participation coefficient at the stimulation site for different runs for each run for each subject. **B**. The best fitting model (see Supplementary Table 1). **C**. Posterior distributions of the intercept and the slopes associated with median ΔPC_within_, median ΔPC_outside_, and max Δz_outside_. The grey lines indicate the 90% credible region, the black square is the median and the dots indicate the min/max value. **D**. The median ΔPC_within_ plotted against the PC of the stimulation site. **E**. The median ΔPC_outside_ plotted against the PC of the stimulation site. **F**. The max Δz_outside_plotted against the PC of the stimulation site. All values in D-F are z-scored as they were standardized before the statistical model was quantified. The solid line draws the slope using the posterior median and dotted lines indicate the possible slopes from the possible range of slopes within the 90% credible interval.

We found that the topological configuration of the network following stimulation was dependent on the PC of the stimulation site. Inspecting the best fitting model (Figure 5BC), we found a negative relationship between the stimulation site’s PC and median ΔPC_within_ (Figure 5CD):when stimulating a node with less between-community connectivity, there is an increase in stimulation site’s community integration following stimulation (and *v*.*v*.). The median ΔPC_within_ posterior distribution had a median of −0.46 (99.6% of the posterior distribution was below 0 and the 90% credible interval between −0.74 and −0.18; Figure 5CD). Conversely, the opposite was found for the median ΔPC_outside_ and stimulation site’s PC. Interestingly, we found that if the stimulation site had a higher PC, then communities other than the stimulation site saw an increase in integration. The posterior distribution here had a median at 0.29 (95.7% of the posterior distribution was above 0, and the 90% credible interval between 0.01 and 0.57; Figure 5CE). Finally, aside from increased integration, the node with the largest z outside the stimulation site’s community also increased its within-community connectivity when the PC at the stimulation site was high (the posterior distribution median at 0.29, 97.6% of the posterior distribution was above 0, and the 90% credible interval between 0.05 and 0.54; Figure 5CF). In summary, the results of RQ2 showed that, if a stimulation site has more connectivity with other communities, it affects the median PC of those communities (and *v*.*v*.).With high PC stimulation sites, stimulation was also shown to subsequently affect the within-community connection for provincial hubs in other communities within the network. This reveals that the topological role of the stimulation site is of importance for what type of widespread network changes that occur following electrical stimulation, and more generally confirms the predictions of functional network organization based on resting fMRI network analyses.

### Additional post-hoc analyses

We considered whether there was any difference between cortical and subcortical stimulation sites (Fig. 4A). We qualitatively contrasted the 8 summary measures. There was a weak trend suggesting an increase in median ΔPC_outside_ for subcortical nodes. However, more data would be required in order to arbitrate whether such a difference exists (See Supplementary Figure 4).

## Discussion

This study capitalized on a unique dataset in order to answer pre-registered hypotheses about the brain’s mesoscale functional network organization. First, we found that the stimulation site’s community generally increased its participation coefficient following stimulation. This shows that stimulating a node generally pushes its entire community towards heightened integration with other brain networks. Second, we show that the topological role of the stimulation site moderates the degree to which stimulation results in changes in integration. If a stimulated node has lower participation (i.e., fewer connections with other communities), then the increase in participation is typically more confined to the stimulation site’s community. Conversely, if a stimulated node has high participation, and thus has more diverse connections to other communities, there was an increase in the integration of nodes outside the stimulation site’s community. Finally, the largest provincial hub outside of the stimulation site’s community also increased its within-community connectivity if stimulating node’s with high participation. Taken together, these results confirm that intracranial electrical stimulation heightens meso-scale network integration as a function of topology of the site that was stimulated.

Our results advance our understanding of the effects of brain stimulation by placing neural stimulation within the context of the global brain network. Previous work has uncovered many important insights about features relatively local to the site of stimulation. For instance, direct stimulation leads to increased local excitability of neurons that could be predicted from pre-stimulation connectivity profiles (Keller et al., 2018) that in turn have been linked to changes in local BOLD responses (Tolias et al., 2005). Transcranial magnetic stimulation (TMS) has been considered to effect system-level changes (Tang et al., 2017), but BOLD studies coupled with TMS show inconsistent changes in connectivity following stimulation (Eldaief et al., 2011; Rounis et al., 2006; Wang et al., 2014), making it hard to determine general principles from these studies. Further, optogenetic work has demonstrated that increases in functional connectivity can persist after stimulation (Yazdan-Shahmorad et al., 2018), but has not used the vantage point to quantify topological effects. Our work complements these findings by adding: (1) the whole-brain coverage provided with concurrent fMRI; and (2) novel network-level analyses to quantify the widespread reconfiguration of nodes caused by the intervention of electrical stimulation. Another major difference between our work and previous work is that we directly measured the network change that occurs during phasic, ongoing stimulation, whereas previous work has largely focused on the persistent effects of stimulation. This enabled us to look at instantaneous changes at the network level that would be impossible to investigate otherwise.

The results of this study suggest that the topological signature of individual regions acts to constrain their activity. Specifically, we found that the topological signature of a region prior to stimulation was associated with the manner in which the functional network signature of the brain was able to reconfigure following the injection of electric current. While not tested explicitly in this study, the implication for this result is that the topological signature of the brain could provide important constraints on the manner in which different neural regions could respond across unique cognitive contexts. Previous research has found a wealth of evidence regarding the importance of states of more integration and/or segregation while performing cognitive tasks (Bassett et al., 2015; Cohen & D’Esposito, 2016; Cole et al., 2013; Fransson et al., 2018; Hearne et al., 2017; Shine et al., 2016). Increased excitability of local stimulation has a complex effect on the topological reconfiguration of the network. This work determined a number of factors that help determine what kind of reconfiguration occurs following the stimulation, however the extent that these reconfigurations would influence cognition or behaviour remains unclear but could be formally tested using es-fMRI if combined with a task. Further extensions of this work could consider which additional factors help determine the topological reconfigurations following electrical stimulation. Two possible candidates are (1) the specific connectivity state when stimulation occurs and (2) whether there are time-varying effects that show the spread of the topological change.

A potential shortcoming of our approach is that our analysis focussed entirely on network-level hypotheses, with little regard for the specific brain regions being stimulated. This limitation means that, throughout the analyses, we have been neutral to which community the stimulation site comes from, which is an idealization of the organization of the brain. For instance, while our approach does afford us the ability to consider the general effects of electrical stimulation, it’s possible that different brain networks may have more inherent topological variation, which cannot be detected here. Secondly, the choice of stimulation site was entirely clinically driven, and hence did not represent an unbiased coverage of the whole brain, however this problem can be solved by future studies on an *ad hoc* basis. Another possible cause for concern is the nonuniform distribution of runs per subject (i.e., some subjects contributed multiple runs, others fewer) in the final analysis pooling a total of 41 runs. This trade-off was necessary given the noise in the data, our requirement of knowing the stimulation site’s community, and insufficient runs in order to hierarchically model the effect. A final limitation is that we noted seven deviations or changes from the preregistration. At times, this was due to some choices not having been explicitly stated in the preregistration. Other deviations, most notably how the stimulation sites were assigned a community and the model used in analysis 2, were due to discovering that the specified analyses were not feasible given the data. See the section *Explicit deviations from preregistration* in the supplementary materials for a complete list of deviations and motivations behind the deviations.

## Conclusion

Here we provide evidence of causal reconfiguration of network properties following intracranial stimulation in humans. Our results demonstrate specific network-level effects of electrical stimulation on the brain that were dependent upon the topological signature of the stimulation region. In doing so, we provide a crucial, systems-level context for previous work that helps to visualize the brains’ response to invasive stimulation at the whole-brain level. Our results suggest that future work investigating different stimulation strategies (such as TMS and tDCS), both in healthy and clinical cohorts, may potentially benefit from a similar topological viewpoint.

## Methods

The project is a preregistered study (https://osf.io/pdhfu/). There were several deviations that are listed in the “deviations from preregistration”.

### Data

The data consisted of 26 subjects, two sessions (*preop* and *postop*). In the preop session, there was a resting state task with regular BOLD fMRI and a T1-weighted (T1w) anatomical scan. In the postop session, there were multiple runs of es-fMRI (number of runs differed per subject). The es-fMRI runs contained no explicit task for the subject. There were periods of repeated intracranial electrical stimulation for 30 seconds at a stimulation site while simultaneously recording BOLD. The dataset is available on *OpenNeuro*.*org* (Accession Number: ds002799) and more information about the dataset is available in the data descriptor article (Thompson et al., 2020).

### Preprocessing

Data were prepared for statistical modeling using fMRIPrep v1.5.1RC Esteban et al., 2019). All the details regarding the preprocessing step with fMRIPrep are reported in Thompson et al 2020 for the dataset. After minimal preprocessing, a modified version of fMRIdenoise (v0.1.1, customized) (https://github.com/wiheto/fmridenoise/tree/9a8587) was run to evaluate which denoising pipeline was considered optimal. The pipeline ‘24HMP_aCompCor_SpikeReg_4GSR’ was chosen, which removes 24 movement regressors, aCompCor, spike regression, and four global signal regressors. Further, fMRIDenoise applied a band-pass filter that was applied at this step (0.008 − 0.1 Hz).

As a starting parcellation, we used 400 regions of interest from the cortex (Schaefer et al., 2018), ten cerebellar ROIs (King et al., 2019), three amygdala ROIs (Tyszka & Pauli, 2016) and the Harvard Oxford subcortical atlas (Desikan et al., 2006; Frazier et al., 2005; Goldstein et al., 2007; Makris et al., 2006) with the amygdala region removed. All atlases were converted into the same MNI space (TemplateFlow ID: *MNI152NLin2009cAsym*). The motivation behind using multiple amygdalae ROIs was due to many of the stimulation sites being there. If any overlapping voxels existed between the different atlases, which sometimes occurred at the boundaries, these voxels were dropped.

There were known susceptibility-derived distortions in the data (see Thompson et al., (2020)) exacerbated by the tissue-implant interfaces, which include nonlinear spatial mislocalization of signal and signal dropout. As a result, some regions of BOLD datasets do not reach the location accuracy and/or signal-to-noise ratio required for analysis. To maximize the number of voxels used in each subject, we created *included voxel* masks for each session (*preop* and *postop*) as follows. We concatenated the average BOLD intensity per voxel. We reasoned that noisy voxels would congregate in the lower tail of the voxel intensity distribution (see Supplementary Figure 2 that compares preop and postop distributions). We then fitted a Gaussian mixed model (scikit-learn 0.21.3 (Pedregosa et al., 2011)). The number of Gaussians ranged between 1-5. We then visually inspected the classification of Gaussians along with the distribution of voxel intensity (see Supplementary Figure 2 for an example). We decided upon using the model with four Gaussians and dropped all voxels classified to belong to the mixture component with the lowest intensity. Two of the co-authors (WHT, OE) reached the decision together through visual inspection of the data. Next, we created parcellations for each subject (one for each preop and postop sessions). Within each parcel, we evaluated whether most of its voxels were still present or excluded after the previous step. If a parcel had a voxel survival rate lower than 50%, then it was removed for that subject’s parcellation. Otherwise, a time series for each parcel was extracted by averaging across all surviving voxels. Time series extraction was done using nilearn v0.6.0a0 (Abraham et al., 2014). See Supplementary Figure 3 for an example of the parcellation overlayed on the data.

### Community detection

The aim of the community detection step was to derive unique subject-specific communities for the custom-built parcellation but with a resolution of a standard brain network template. To achieve this, all postop runs for each subject were concatenated. Functional connectivity was determined using Pearson correlations for all the ROIs. The communities were derived using the Leiden algorithm (Traag et al., 2019) using the (Reichardt & Bornholdt, 2006) null model implemented in iGraph for python v0.7 (Csárdi & Nepusz, 2006). In order to set the resolution parameter, we aimed to maximize the overlap with a standardized cortical template (the Yeo 7 template (Yeo et al., 2011)). The resolution parameter ranged between 0.5 and 2.5 in steps of 0.01. For each resolution parameter, the algorithm ran until convergence (i.e., no improvement). We determined the best fitting resolution parameter with the adjusted mutual information (AMI) in scikit-learn 0.23.3 (Pedregosa et al., 2011) between the remaining cortical nodes and the corresponding nodes for the Yeo et al. 7 network template. We chose the resolution parameter by taking the largest AMI score for each subject.

Each stimulation site was assigned to a community. We identified the stimulation site’s coordinates to be equidistant between the two stimulating channels (one channel got the leading positive phase of stimulation). We then placed a 6mm sphere surrounding this spot. We then checked which parcels overlapped the most with the sphere. This parcel’s community became the community of the stimulation site. If the parcel associated with the stimulation site was to be excluded or was a singleton or 2-node community, then that run was excluded from all subsequent analyses that required there to be a community for the stimulation site (see below). The motivation for singleton/2-node community exclusion was that the module degree z-score would always be zero for such subjects.

### Subject inclusion

The inclusion of data was done in multiple steps. First, screening of fMRIPrep’s summary reports to evaluate the preprocessing of each run. Second, by quantifying if the average framewise displacement was greater than 0.5 to remove runs that had excessive head motion. Third, we only used runs that had a community associated with the stimulus site and that community was not a singleton or 2-node community. The first two exclusion steps dropped subjects and runs from the entire analysis respectively. The final step only excluded runs in steps that required a stimulation site. In the community detection phase, for example, where no stimulation site was needed, these runs were included.

### Network measures

For each node, the participation coefficient (PC) and the module degree z-score (z) were calculated using bctpy v0.5. The PC was calculated on positive edges only. PC measures the proportion of an edge’s weights that are outside of its community. z measures the z-scored strength of a node within its community. Together they depict information about a node’s behaviour as a connector hub (high PC, connecting communities) and a provincial hub (high z, connecting nodes within a community). See (Guimera et al., 2005) for more information about these measures.

Next, in order to calculate the effect of the stimulation, we calculated the difference between the stimulation, namely:

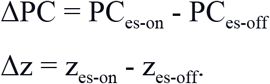

The first five seconds of each es-on and es-off period were discarded to avoid any spill-over effects due to the sluggishness of the BOLD signal.

In order to summarize the behaviour of the stimulation site’s community and outside its community, we derived four different metrics for both ΔPC and Δz. These metrics were: (1) median within the stimulation site’s community, median difference in PC (or z) (abbreviated to median ΔPC_within_ and median Δz_within_), (2) outside the stimulation site’s community, median difference in PC (or z) (abbreviated to median ΔPC_outside_ and median Δz_outside_), (3) within the stimulation site’s community, the max difference in PC (or z) (abbreviated to max ΔPC_within_ and max Δz_within_),, (4) outside the stimulation site’s community, the max difference in PC (or z) (abbreviated to max ΔPC_outside_ and max Δz_outside_),.

### Statistical model

In the first analysis, we tested whether there was a difference between the network metrics within and outside the stimulation site’s community. We tested whether the differences between es-on and es-off for both PC and z were more substantial in the stimulation community compared to outside the stimulation community. The null hypothesis was that there would be no difference. We performed four statistical tests, comparing: max PC, max z, median PC, and median z. The p-values were Bonferroni corrected (0.05). 10,000 permutations where each run’s within or outside community membership were randomly permuted to calculate the null-distribution.

In the second analysis, we constructed Bayesian models using pymc3 (v3.7) (Salvatier et al., 2016). The models all had the stimulation site’s participation coefficient as the dependent variable. The independent variable(s) were some combination of the 8 summary statistics of the difference between es-on and es-off. An intercept was modelled as well. All variables were standardized beforehand by removing the mean and dividing by the standard deviation. All different permutations of the dependent variables were run as well as one control model with just an intercept. The model was specified as follows:

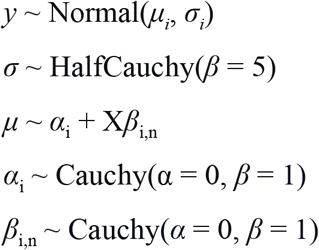

The priors were all weakly informed. The size of X and *n* in *β*_i,n_ depend on the number of independent variables in that particular model. For each model, two Markov chain Monte Carlo (MCMC) chains were sampled 10,000 times after 1,000 tuning samples. LOO and the widely applicable information criterion (WAIC, a generalized version of the Akaike information criterion, AIC) were used to evaluate the models (Supplementary Table 1 & 2). However, based on the warnings issued during the WAIC model fit, it was not used to choose the model (but the result was the same). The chosen model from the LOO had the median change in median ΔPC_within_,median ΔPC_outside_, max Δz_outside_ as the independent variables. The chosen model was deemed sufficient by visual inspection of the MCMC fit, the Gelman Rubin statistic being close to 1 for all variables, and posterior checks for the mean and IQR value being within the acceptable range of 0.2-0.8 (mean: p=0.4959, IQR: p=0.7794).

### Code for analysis

The entire code for the project can be found at https://github.com/wiheto/esfmri_connectivity under a Apache-2.0 License. Further, this repository has been linked to the preregistration at OSF, and all pull requests can be tracked seeing all the changes made.

## Supplementary Figures

**Supplementary Figure 1.**
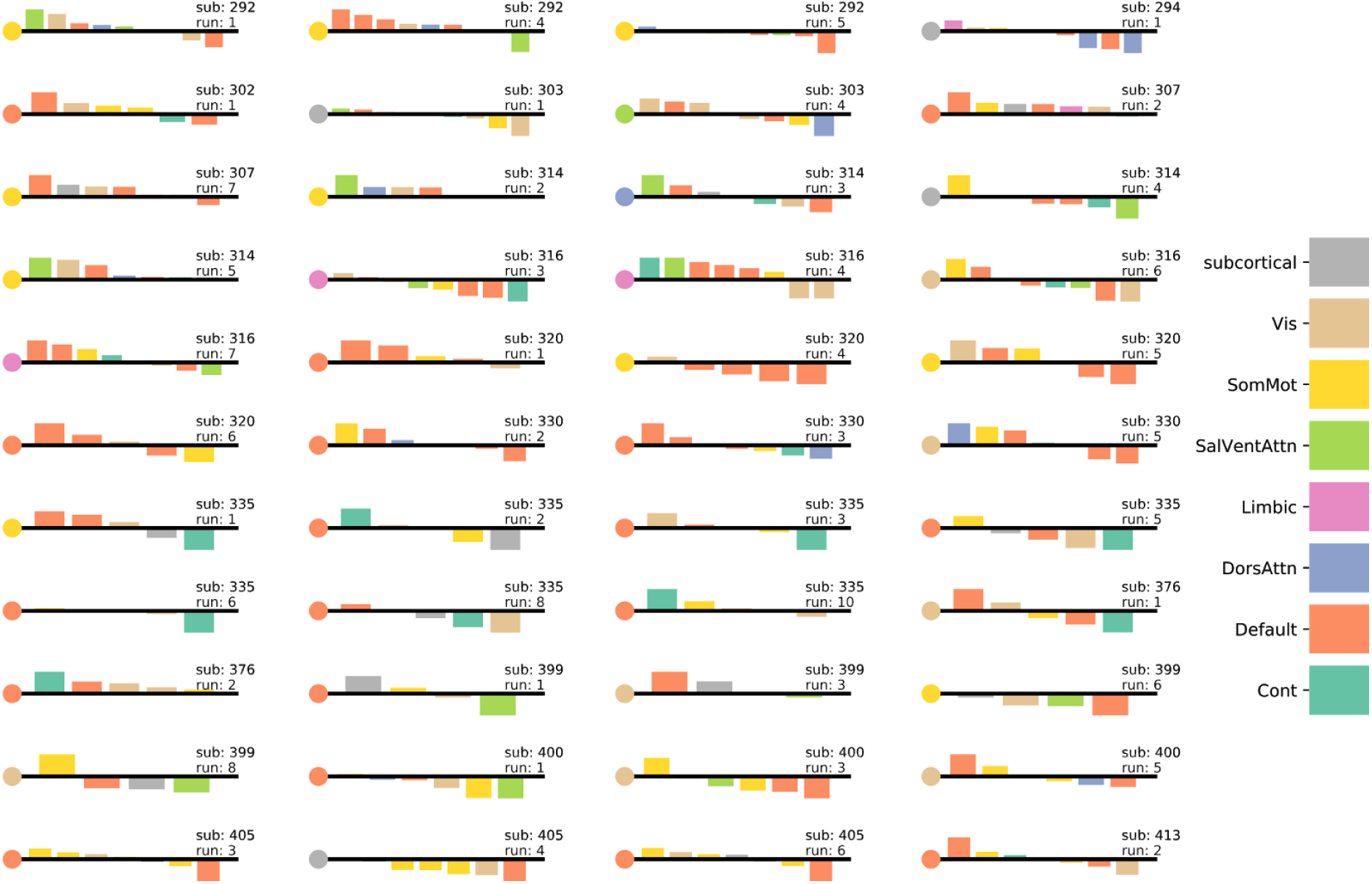
The change in median between-community strength with the stimulation site had diverse profiles between subjects. Change in median community strength with the stimulation site (marked by the circle). The specific y-values are arbitrary here as the contrast is the profile of the different bars (i.e. if all bars behave the same or if there is greater variance). Bars are ordered from highest to lower. Only non-singleton communities are shown. The black line always indicates zero. Community labels are assigned by finding the largest number of nodes the communities overlap with the Yeo 7 template, which can lead to the same community appearing multiple times.

**Supplementary Figure 2.**
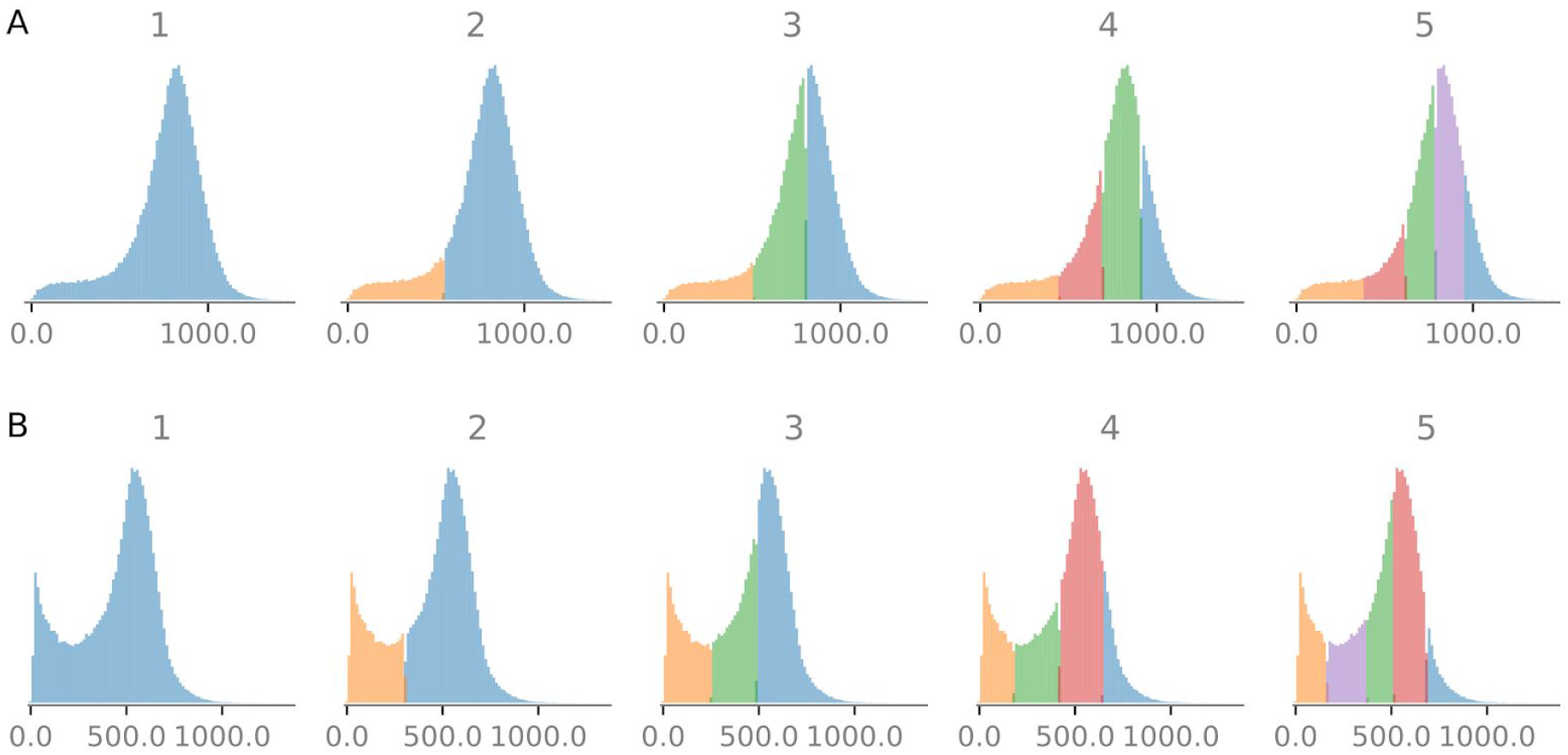
Example of average voxel intensity between (A) preop (resting) (B) and postop (electrical stimulation) sessions for a single subject. The figure shows how the distributions of voxels were classified by varying the number of Gaussian distributions. The electrical stimulation trials have a large tail close to 0. We chose 4 Gaussian distributions for all subjects and classed all voxels belonging to the lowest Gaussian as bad voxels.

**Supplementary Figure 3.**
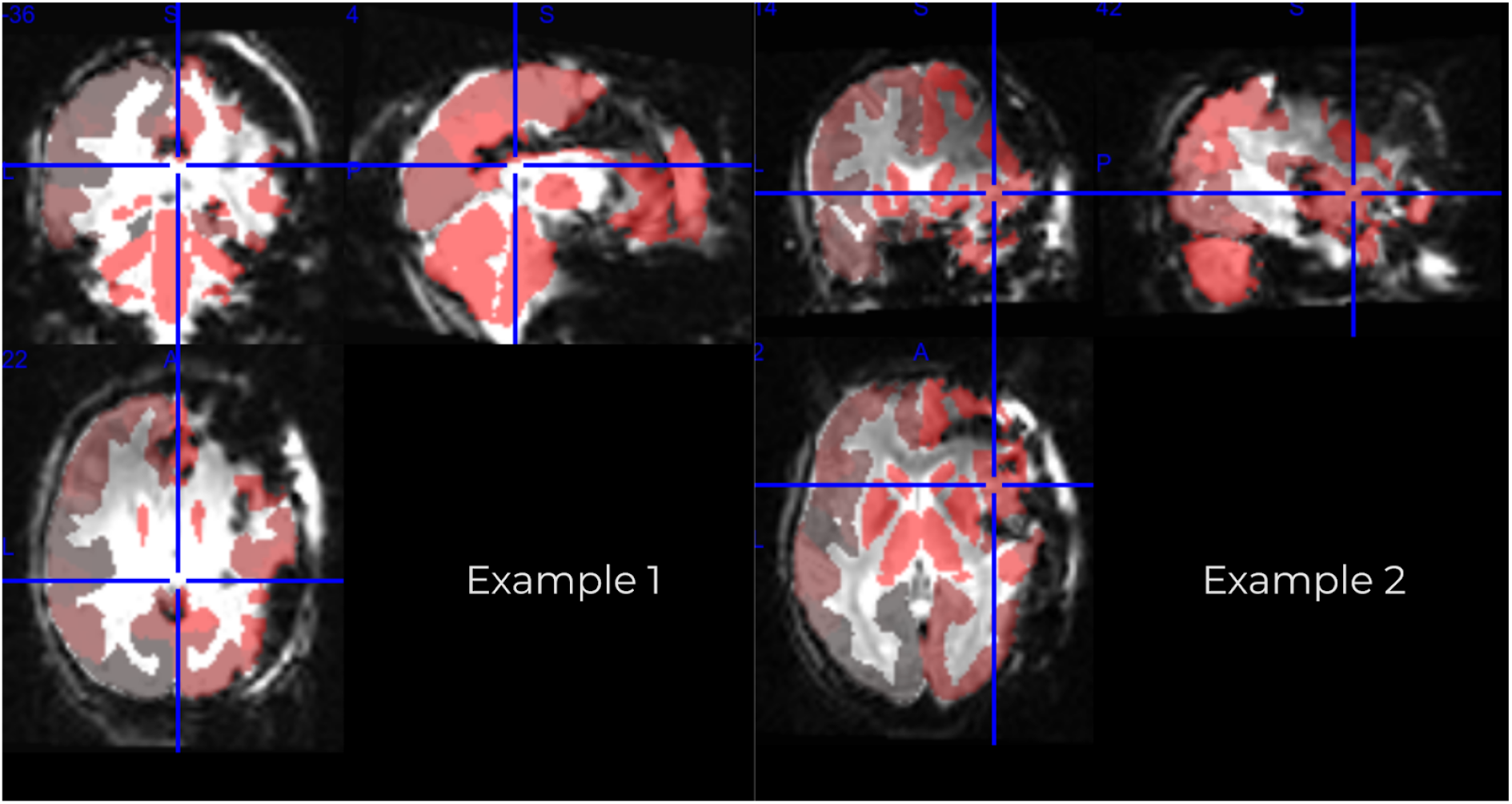
Example of parcellation overlayed over subject. Two examples of the parcellation overlayed on a single volume of the BOLD data during electrical stimulation. The different shades of red represent different parcels in the parcellation when using only the good voxels. Here we see that the procedure to remove bad voxels and parcels has done a good job at avoiding the considerable noise which exists for each subject.

**Supplementary Figure 4:**
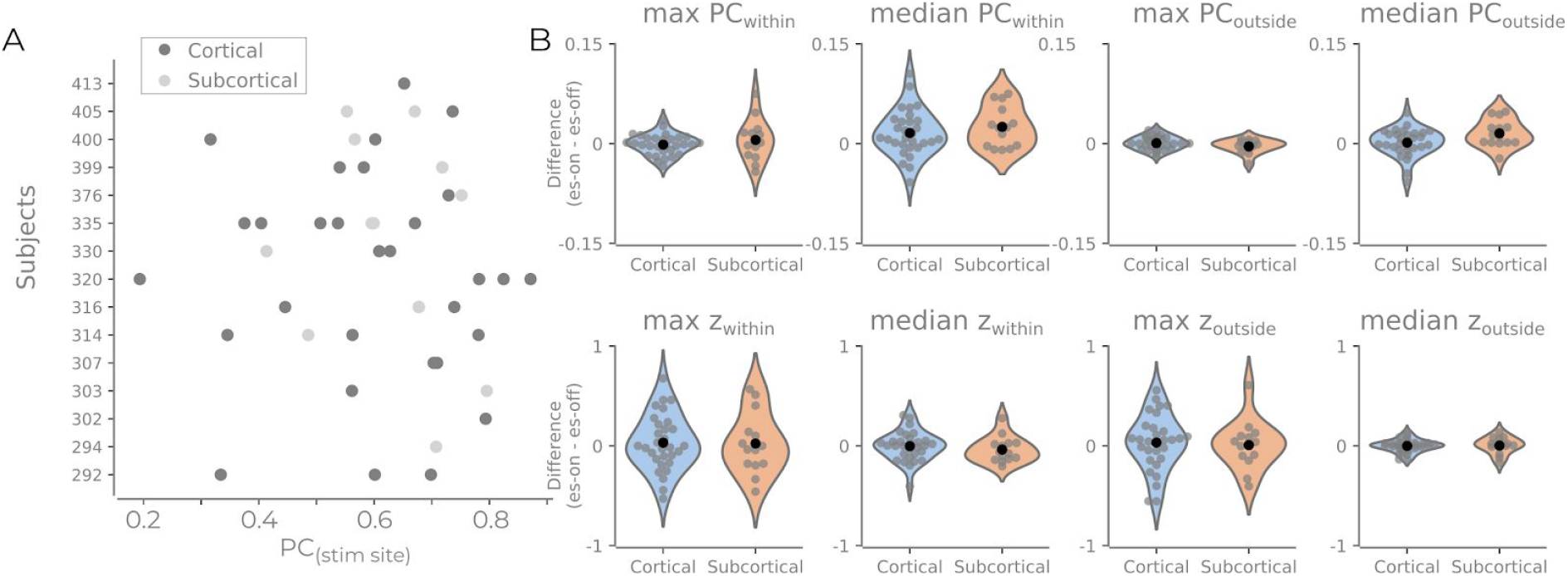
Difference in the eight different difference summary measures for subcortical and cortical stimulation sites. As part of another *post hoc* analysis, we asked if there was any difference between cortical and subcortical stimulation nodes. A. Same as Figure 5A but marking the cortical and subcortical nodes.. It is possible that subcortical nodes affect the network in a qualitatively different way. B. We descriptively checked all eight summary measures. No clear noticeable difference was found. There is a trend of an increase in median ΔPC_outside_ for subcortical nodes, however, the trend was weak and more data would be required in order to arbitrate such a relationship.

**Supplementary Table 1: seperate excel sheet**

**Supplementary Table 1**. LOO scores for each of the different model combinations. The models show the different summary delta variables used as independent variables in a model with the stimulation site’s participation coefficient as the dependent variable.

**Supplementary Table 2: seperate excel sheet**

**Supplementary Table 2**: LOO scores for each of the different model combinations. The models show the different summary delta variables used as independent variables in a model with the stimulation site’s participation coefficient as the dependent variable. Note that all model rows are flagged with a warning, which indicates that WAIC may be an unreliable evaluation of the model fit.

## Supplementary Text

### Explicit deviations from preregistration

1. Selecting the best de-noising pipeline was not apparent from the different metrics as we had hoped in the preregistration. The choice (24HMP_aCompCor_SpikeReg_4GSR) was thus informed by which pipeline is the most rigorous, appears to correlate the least with the null model, and by looking at the connectivity matrix in the fMRIDenoise reports.
2. The procedure to threshold bad voxels with GMM, we chose to keep all but the lowest GMM instead of only keeping the highest. This is a more sound method than specified in the preregistration as it allowed for removing the worse component and keeping the majority of the data. This decision was based on the visual inspection where it was determined that only keeping the highest gaussian would remove too much data. This can be seen in Supplementary Figure 2 where the distributions of voxel intensity is comparable between preop and postop for all but the lowest Gaussian. This choice was made before continuing with the data analysis.
3. Adjusted mutual information is used instead of normalized mutual information with regards to comparing the cortex communities with the Yeo 7 template communities. Otherwise, this produced one large community or very small communities as having the best NMI. AMI corrects for this.
4. Only positive edges are considered in the community detection step and the participation coefficient. This was not explicitly stated in the preregistration.
5. There were some runs where there were multiple simultaneous stimulation sites during the es run. These are excluded. This was not explicitly stated in the preregistration, but without doing this would yield a different type of comparison than is done for all other runs. These were excluded before continuing with the analysis.
6. It was unclear in the preregistration how the community detection in analysis one was to be calculated and how a community was to be assigned to the stimulation site. Instead of using the preop data (which would reduce the number of participants), we decided to concatenate all the es-runs and use this to identify the stimulation sites community and categorize the topological role of the stimulation sites. We calculated the FC on the concatenated FC data per subject and derived the communities from this. To identify the stimulation site, we placed a sphere between the two channels and identified the stimulation site’s parcel as the parcel that had the most overlap with the sphere. The preregistration stated that this would be 3mm, but for some subjects, no parcel lay in that radius. Subsequently, we expanded the radius of the centre by 1mm increments until all subjects had a parcel associated with the stimulation sites. All subjects had a parcel that could be assigned at 6mm radius, so this was the choice. This choice was made to maximize the amount of data in the next step. Importantly the overlapping ROIs were calculated on the parcellation template, not the subject maps. Thus, there was still a possibility that the overlapping parcel was not included in the subject’s mask. If the parcel identified as the stimulation site was not in the subject mask, then the run was excluded.
7. The initial pre-registered plan was to calculate the connectivity per stimulation sequence (i.e. multiple connectivity estimates per run) and then place this in a hierarchical statistical model with the subject as a layer. When writing the preregistration, we did not realize that this would only be <10 data points per stimulation run. We deemed this too few to get accurate estimates, and this analysis was never performed. Instead, we calculated the estimates for each run (as the first analysis). Because of this change, we also needed to use the difference (i.e. es-on - es-off) instead of just es-on. Without comparing the change to baseline, the values will be hard to compare, as the stimulation sites belong to communities with different PC-properties. Thus, we had to use the displacement values here to achieve more meaningful comparisons. The preregistration also only specified the median values would be used, but left open the possibility of including the max values as well. We saw no reason to not also include the max values in the model evaluation.

